# Combinatorial screening of nanoparticles for nose-to-brain RNA delivery to modulate neuroinflammation after traumatic brain injury

**DOI:** 10.64898/2026.05.27.727842

**Authors:** Yihan Xing, Vy Do, Zunkai Xu, Clara Do, Canyuan Yang, Ziyue Zhu, Jingjing Gao

## Abstract

Traumatic brain injury (TBI)-induced neuroinflammation can evolve over weeks or months, contributing to ongoing secondary damage and worsening neurological recovery. RNA-based therapeutics hold great potential to regulate inflammatory signaling, but delivery to the brain remains challenging because of the blood brain barrier. Intranasal administration offers a direct non-invasive route for brain access but is often limited by low delivery efficiency. Here we constructed a combinatorial library of DNA-barcoded 103 lipid nanoparticles by varying lipid components with diverse headgroup chemistries and bioreactive moieties. A high throughput screening of these nanoparticles *in vivo* following intranasal administration led to the identification of top candidates with highest brain accumulation. Intranasal delivery of an antagomir targeting microRNA-9-5p loaded nanoparticle effectively reduced cerebral microRNA-9-5p levels, suppressed inflammatory markers, and improved neurological outcomes in TBI. These results demonstrate a systematic approach for optimizing intranasal lipid nanoparticle design and support the feasibility of RNA delivery to modulate neuroinflammation after brain injury.

## Introduction

Traumatic brain injury (TBI) represents a paramount global health challenge, affecting over 50 million individuals annually and incurring economic costs exceeding $400 billion^1^. Moreover, TBI is projected to remain a leading cause of neurological disability through 2030^2^. Beyond the immediate mechanical insult, TBI rapidly initiates a secondary injury cascade dominated by neuroinflammation, characterized by early microglial activation, immune cell infiltration, and evolving neuronal damage^3^. Although these processes begin within hours after injury, they can persist and evolve over days to weeks, progressively reshaping the neuroimmune microenvironment^4^. Critically, if early inflammatory amplification is not effectively restrained, it may transition into sustained or chronic neuroinflammation, contributing to ongoing neuronal loss and long-term neurological deficits^5^. Therefore, timely modulation of post-injury neuroinflammation is essential not only to reduce acute damage but also to prevent the establishment of maladaptive chronic inflammatory states.

Although nucleic acid therapeutics hold promise for intercepting this cascade, their systemic utility is severely restricted by the blood-brain barrier (BBB)^6, 7^. Physical breaching of the BBB after TBI provides a narrow window for systemic delivery, but an important limitation is that secondary injury after TBI can persist for months and require repeated dosing well beyond this transient period^8, 9, 10^. Moreover, systemically administered nucleic acid therapeutics, particularly LNP formulations, often accumulate in peripheral organs due to lipid metabolism and hepatic uptake, thereby limiting effective brain targeting^11^. Consequently, there is an urgent need for non-invasive strategies that bypass the BBB and deposit therapeutic payloads directly into the injured brain. Intranasal (i.n.) administration offers a promising strategy to bypass the blood-brain barrier, providing direct access to the central nervous system *via* the olfactory and trigeminal nerve pathways^12, 13, 14^. However, the therapeutic potential of this route is fundamentally restricted by rapid mucociliary clearance^15, 16^. This physiological defense mechanism typically eliminates deposited particulates within 15 to 20 minutes, significantly shortening the window for drug absorption^17^. Therefore, extending effective nasal residence while preserving downstream transport efficiency represents a central challenge for intranasal nanocarrier design.

Previous studies have shown that physicochemical properties of nanoparticles influence their interactions with nasal mucus and epithelial barriers^18, 19^. For example, cationic formulations such as chitosan have been associated with increased mucoadhesion and enhanced cellular uptake *in vitro*^18, 20^, whereas neutral or anionic formulations and surface-shielding modifications such as PEGylation have been reported to facilitate deeper mucus penetration in model systems^16, 19, 21, 22^. However, these properties can exert opposing effects on residence time and transport, and current findings from mostly *in vitro* assays^19, 21, 22^ are not always consistent^23, 24^. It remains unclear which specific feature of the nanoparticles most effectively support nose-to-brain delivery under physiological conditions.

Herein we engineered a combinatorial library comprising 103 lipid nanoparticles that integrates diverse headgroup chemistries and bioreactive moieties to improve nose-to-brain delivery efficiency (Figure 1). Using DNA barcoding coupled with next-generation sequencing, we performed a high-throughput *in vivo* screening to enable unbiased, head-to-head quantification of brain distribution across formulations. This approach allows us to delineate exactly which chemical features drive successful brain delivery and we identified a lead nanoparticle formulation with markedly improved brain delivery efficiency.

**Figure 1.**
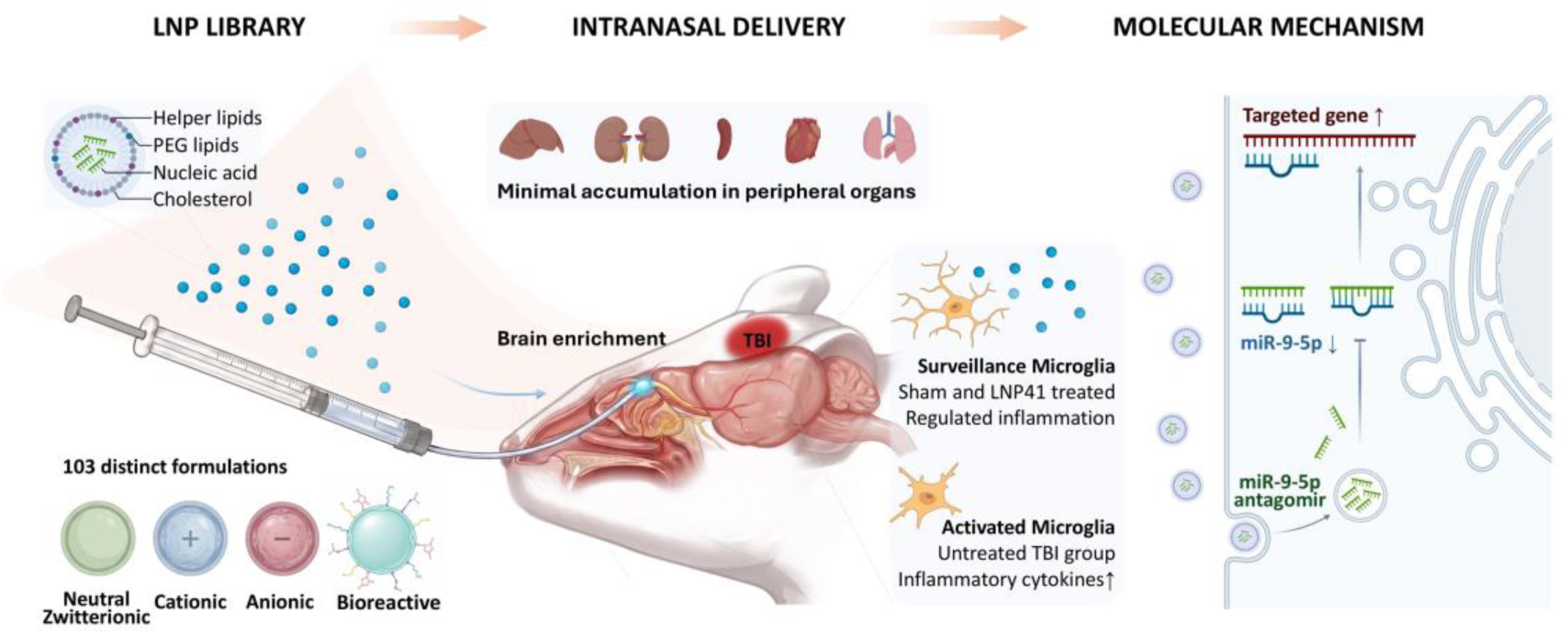
Schematic illustration of intranasal LNP delivery and therapeutic mechanism to attenuate neuroinflammation post TBI. A combinatorial LNP library was constructed by varying helper lipid and PEG lipid composition to enable systematic screening of brain-targeting formulations. Intranasal administration facilitated direct nose-to-brain delivery, resulting in preferential accumulation in the brain with minimal distribution to peripheral organs. In the untreated traumatic brain injury (TBI) condition, microglia exhibit an activated phenotype characterized by elevated miR-9-5p expression and increased pro-inflammatory cytokine production. In contrast, delivery of antagomir-miR-9-5p via the lead LNP formulation (LNP41) modulates microglial activation toward a surveillance-like state, accompanied by reduced inflammatory signaling. At the molecular level, miR-9-5p inhibition regulates downstream gene expression, contributing to attenuation of neuroinflammation.

To demonstrate the therapeutic potential, we next used the lead LNP formulation to deliver an antagomir against miR-9-5p, a key regulator of microglial activation^25, 26^ that is markedly upregulated after traumatic brain injury^26, 27, 28^. miR-9-5p is implicated in modulating pro-inflammatory microglial responses and it has been linked to NF-κB-associated signaling^29, 30, 31^, and the promotion of pro-inflammatory polarization^27, 32, 33, 34^. Given the dynamic spatiotemporal regulation of miR-9-5p after TBI and its contribution to amplification of secondary neuroinflammation, therapeutic suppression during the evolving injury phase represents a mechanistically grounded strategy^25^. In a mouse model of traumatic brain injury, we investigated the efficacy of intranasal administration of miR-9-5p antagomir-loaded LNPs. We demonstrate that this approach effectively reduced the levels of miR-9-5p, significantly mitigated neuroinflammation and resulted in neurofunctional improvement. Collectively, these findings establish a robust therapeutic strategy for targeting post-traumatic neuroinflammation and improving recovery after brain injury, highlighting this platform as a translatable approach for nose-to-brain delivery of RNA therapeutics.

## Results

### Construction and characterization of combinatorial LNP library

To systematically screen LNPs for maximized nose-to-brain delivery, we built a combinatorial lipid nanoparticle library that systematically tunes two key features: charge and bioreactivity (Fig. 2). Using SM102 (ionizable lipid) and cholesterol as a constant core, we first varied the helper lipid to span zwitterionic (DSPC, DSPE), cationic (DSTAP), and anionic chemistries (DSPS, DSPE-succinic acid), and lipids bearing reactive functional groups (e.g., -SH and -NHS ester). In parallel, we varied the PEG-lipid component by incorporating PEG-lipids with different terminal charges (DMG-PEG2000, DSPE-PEG2000-NH_2_, DSPE-PEG2000-COOH), as well as PEG-lipids containing reactive handles (e.g., DSPE-PEG2000-maleimide and DSPE-PEG2000-Ald) (Fig. 2A). By systematically combining these helper- and PEG-lipid variants at defined molar ratios, we generated a library of 103 distinct LNP formulations (Supplementary Table 1), enabling direct evaluation of how lipids charge and reactive moieties influence subsequent brain delivery after intranasal administration.

**Figure 2.**
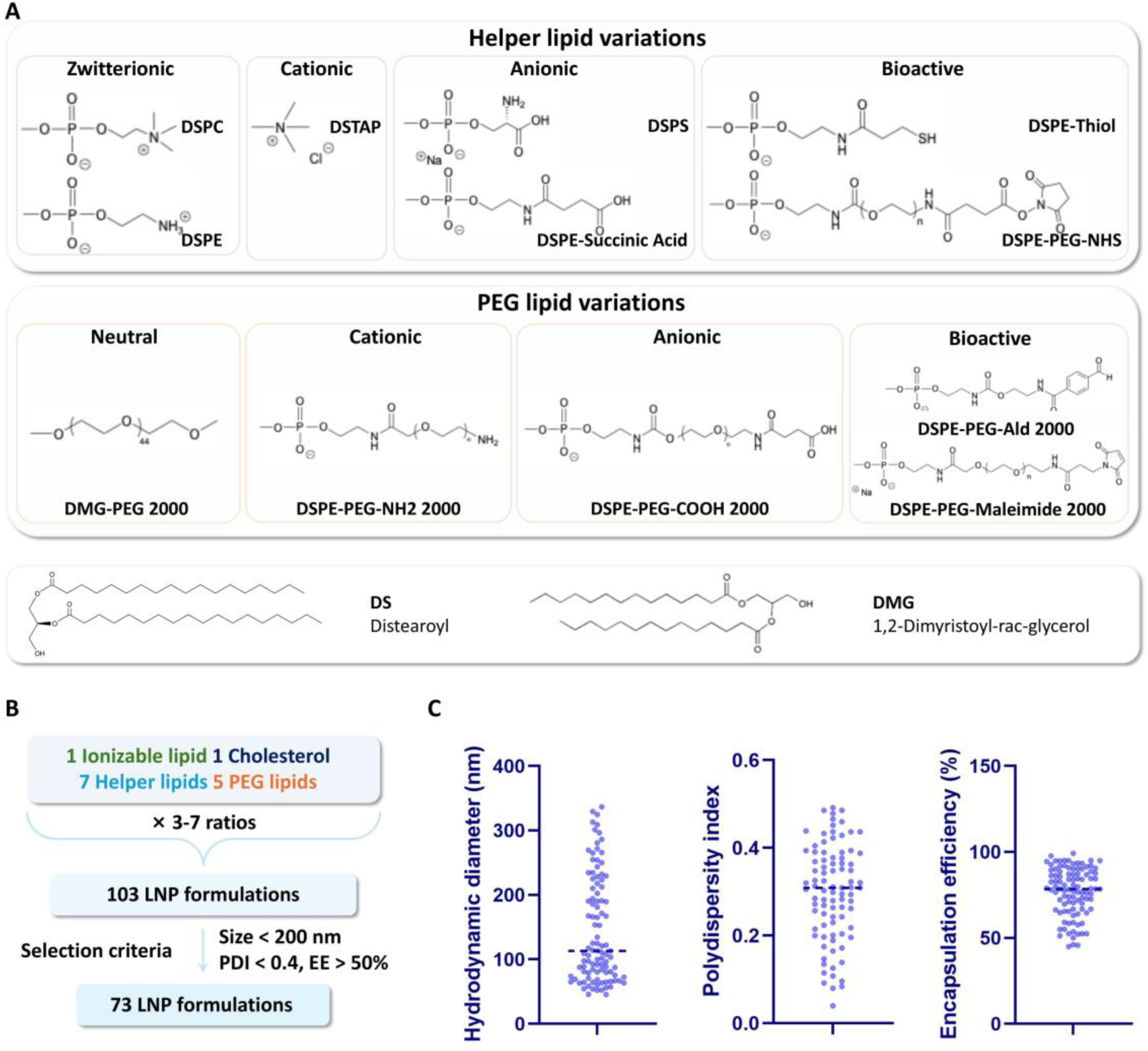
Design and characterization of the LNP library. **A** Helper lipid library classified according to headgroup charge, including neutral, positively charged, and negatively charged lipids. Representative chemical structures of each helper lipid used in this study are shown. PEG lipid library categorized by different terminal charges. In addition to a non-functionalized PEG lipid control, PEG lipids bearing different terminal chemistries were included to modulate LNP properties. **B** Schematic overview of LNP library construction. Each formulation consisted of one ionizable lipid, cholesterol, selected helper lipids, and PEG lipids combined at multiple component ratios, yielding a total of 103 LNP formulations. Formulations meeting predefined physicochemical criteria (size < 200 nm, polydispersity index < 0.4 and encapsulation efficiency > 50%) were retained for subsequent screening. **C** Physicochemical characterization of 103 LNP formulations, including hydrodynamic diameter, polydispersity index, and encapsulation efficiency. Individual data points represent distinct LNP formulations, with box plots indicating distribution across the library.

To enable high-throughput screening *in vivo*, we encapsulate a unique DNA barcode to each chemically distinct formulation and applied stringent quality control filters to the initial library. Formulations with size > 200 nm, polydispersity index (PDI) > 0.4, and encapsulation efficiency (EE) < 50% were excluded. This screening step narrowed the library set to 73 formulations that met the quality control standards and were advanced for *in vivo* evaluation (Fig. 2B). Physicochemical characterization of these 103 formulations showed that most LNPs exhibited hydrodynamic diameters below 150 nm, with a subset extending to ∼300 nm. PDI values were generally between 0.2 and 0.4, while encapsulation efficiencies were robust, largely exceeding 75% (Fig. 2C).

### High-throughput i*n vivo* screening identifies brain-tropic LNPs

To evaluate nose-to-brain delivery in *vivo*, we allocated these 73 formulations into four pooled batches to avoid potential aggregates due to electrostatic attractions or reactions among different functional groups (Supplementary Table 1). Each pooled batch of LNPs was administered intranasally to mice using a catheter-guided approach^12, 35^ (Fig. 3A). This *in vivo* screening workflow allowed for the simultaneous quantification of LNP distribution across different organs (brain, lungs, heart, spleen, kidneys, liver), as well as across specific brain regions and cell types using high-throughput sequencing (Supplementary Fig. C).

**Figure 3.**
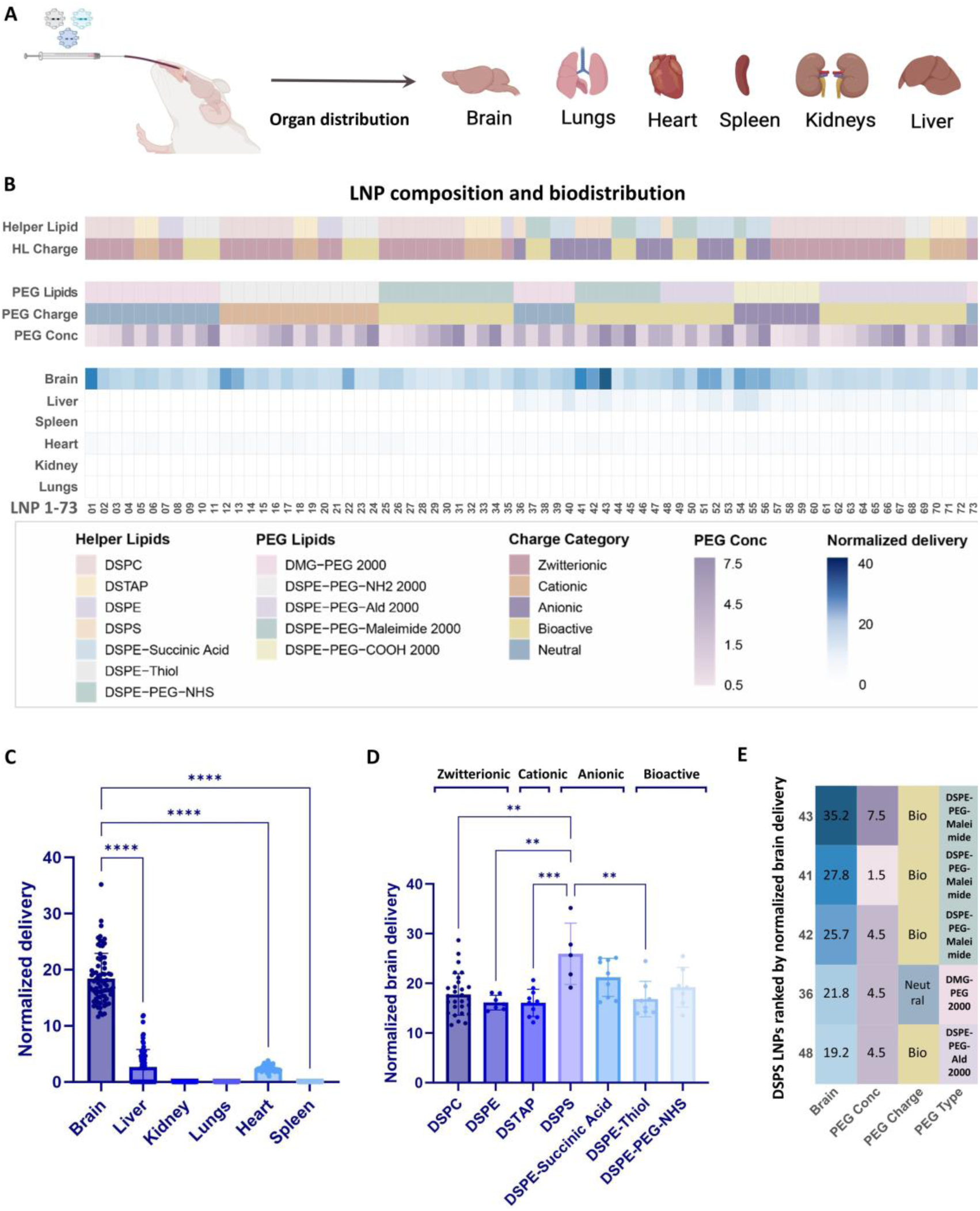
High-throughput *in vivo* screening of intranasal LNP brain distribution. **A** Schematic illustration of intranasal administration of DNA-barcoded LNPs and subsequent assessment of biodistribution across major peripheral organs, brain regions, and neural cell types. **B** Heatmap summarizing LNP library composition and organ-level distribution following intranasal administration. Each column represents an individual LNP formulation, annotated by helper lipids identity, helper lipids (HL) charge, PEGylated lipids identity, PEGylated lipids charge, and PEGylated lipids concentration (Conc). Color intensity indicates normalized delivery across organs. **C** Normalized delivery of DNA barcodes across major organs following intranasal administration, demonstrating preferential enrichment in the brain compared to peripheral tissues. Each dot represents an individual LNP formulation, shown as the mean value across biological replicates. **D** Normalized brain delivery stratified by helper lipid identity. LNPs containing the anionic lipid DSPS show significantly enhanced normalized brain delivery relative to other helper lipid formulations. **E** Heatmap summarizes DSPS-containing LNPs ranked by normalized brain delivery and associated PEG features. Variations in PEG terminal chemistry show modest differences overall, with DSPE-PEG-maleimide contributing to higher normalized brain delivery in specific formulations. Data are presented as mean ± SD. Statistical significance: * *p* < 0.05, ** *p* < 0.01, *** *p* < 0.001, **** *p* < 0.0001.

After normalizing the delivery efficiency based on the barcodes count, we generated a heat map analysis that correlates lipid composition with biodistribution (Fig. 3B). We found that LNP accumulation in peripheral organs such as the liver, kidneys, and spleen was minimal for most formulations, whereas the brain showed significant enrichment (indicated by darker blue intensities) across many formulations. Quantitative analysis of total barcode abundance in each organ further confirmed this brain-targeting capability (Fig. 3C), with the brain accounting for approximately 75% of the recovered relative abundance (Supplementary Fig. D). Together, these results validated the effectiveness of the catheter-based intranasal delivery technique for brain access.

To further examine formulation-dependent effects on brain delivery, normalized brain delivery was firstly analyzed by stratifying LNPs according to helper lipid identity (Fig. 3D). Among the tested formulations, LNPs containing anionic lipid DSPS exhibited significantly higher normalized brain delivery compared to other helper lipids, indicating a strong influence of helper lipid composition on brain delivery efficiency. To further resolve this effect, DSPS-containing formulations were analyzed across different PEG terminal chemistries. Within this subset, DSPE-PEG-maleimide showed the highest normalized brain delivery, whereas other PEG variants exhibited more comparable levels (Fig. 3E).

Analysis across all formulations showed no significant effect of PEG terminal chemistry alone on normalized brain delivery (Supplementary Fig. E). A modest increase was observed for - COOH, while maleimide showed greater variability, including a peak in DSPS-containing formulations, and -NH₂ overlapped with the control. Overall, these results suggest that helper lipid composition plays a dominant role in driving brain delivery, whereas PEG end-group functionality contributes in a more subtle and formulation-dependent manner.

### Stepwise functional screening identifies the optimal LNP formulation

Following the *in vivo* screening, the top 10 LNP candidates with the highest normalized brain delivery were selected for functional validation (Fig. 4A, Table 1). While brain uptake is critical, the ability of LNPs to effectively release their payload and transfect the cells is equally vital. To assess transfection potency, FLuc mRNA was encapsulated into the top 10 formulations and applied to BV2 microglial cells. As shown in Fig. 4B, transfection efficiency varied significantly among the candidates. LNP41 displayed the highest luminescence intensity, followed by LNP12 and LNP01. Consequently, these three formulations (Top 3) advanced to functional *in vitro* screening.

**Figure 4.**
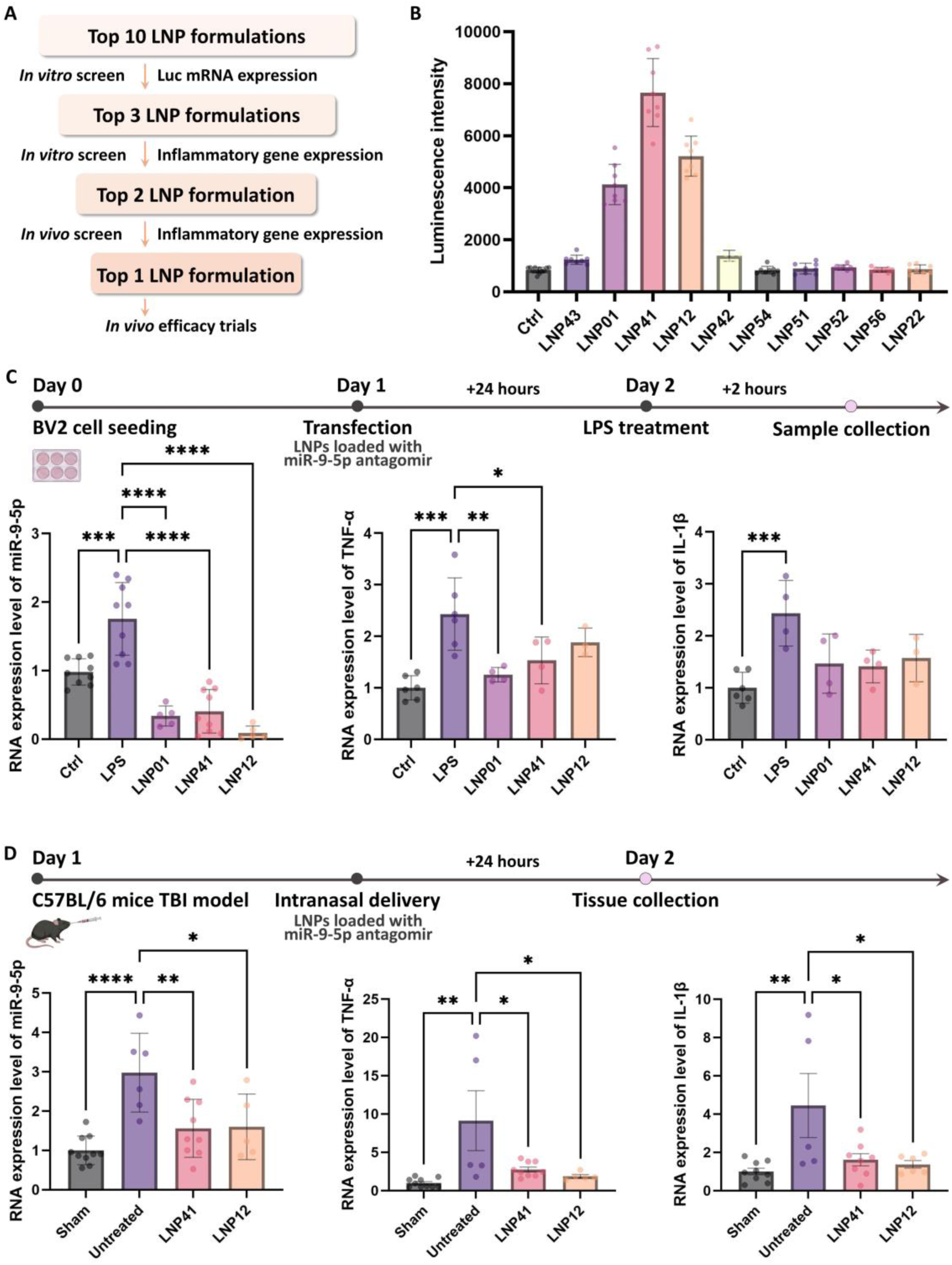
Stepwise functional screening validates the therapeutic potential of lead LNPs *in vitro* and *in vivo*. **A** Schematic workflow of the hierarchical screening process, narrowing down candidates from the Top 10 to a single lead formulation. **B** *In vitro* transfection efficiency of the Top 10 brain-targeting LNPs encapsulating FLuc mRNA in BV2 cells. Luminescence intensity indicates LNP41, LNP12, and LNP01 as the top performers. **C** *In vitro* functional validation of the Top 3 LNPs (01, 41, 12). BV2 cells were transfected with antagomir-miR-9-5p and stimulated with LPS. RT-qPCR quantification shows the relative expression levels of miR-9-5p, TNF-α, and IL-1β. **D** Short-term *in vivo* validation of the Top 2 LNPs (41, 12) in a TBI mouse model. Mice received intranasal LNP treatment post-injury, and brain tissues were analyzed 24 hours. Graphs depict the downregulation of miR-9-5p and pro-inflammatory cytokines (TNF-α, IL-1β). Data are presented as mean ± SD. Statistical significance: * *p* < 0.05, ** *p* < 0.01, *** *p* < 0.001, **** *p* < 0.0001.

**Table 1.** Top 10 brain accumulated LNP candidates.

| No. | LNP ID | Helper Lipid | PEG Lipid | PEG (mol%) |
| --- | --- | --- | --- | --- |
| 1 | 43 | DSPS | DSPE-PEG-Maleimide 2000 | 7.5 |
| 2 | 01 | DSPC | DMG-PEG 2000 | 0.5 |
| 3 | 41 | DSPS | DSPE-PEG-Maleimide 2000 | 1.5 |
| 4 | 12 | DSPC | DSPE-PEG-NH2 2000 | 0.5 |
| 5 | 42 | DSPS | DSPE-PEG-Maleimide 2000 | 4.5 |
| 6 | 54 | DSPE-PEG-NHS | DSPE-PEG-COOH 2000 | 1.5 |
| 7 | 51 | DSPE-Succinic Acid | DSPE-PEG-Ald 2000 | 1.5 |
| 8 | 52 | DSPE-Succinic Acid | DSPE-PEG-Ald 2000 | 4.5 |
| 9 | 55 | DSPE-Succinic Acid | DSPE-PEG-COOH 2000 | 4.5 |
| 10 | 56 | DSPE-Succinic Acid | DSPE-PEG-COOH 2000 | 7.5 |

To evaluate the therapeutic potential of the top LNP candidates in attenuating neuroinflammation, we first examined their ability to deliver antagomir-miR-9-5p in vitro using BV2 microglial cells. miR-9-5p is chosen as the target because it’s a key regulator of microglial polarization^36^, and is markedly upregulated in chronic phase after traumatic brain injury^25^. BV2 cells were transfected with the Top 3 LNPs loaded with antagomir-miR-9-5p, followed by LPS stimulation to induce a pro-inflammatory environment. RT-qPCR analysis revealed that the LPS-only group (LPS) exhibited elevated levels of miR-9-5p and pro-inflammatory cytokines compared to the control group (Ctrl). Treatment with all three candidate LNPs significantly downregulated miR-9-5p expression (Fig. 4C). Importantly, this gene silencing was accompanied by a potent anti-inflammatory effect, evidenced by significant reductions in TNF-α and IL-1β mRNA levels. Among the tested candidates, LNP41 and LNP12 demonstrated the most robust suppression of inflammatory markers. Therefore, they were selected as the Top 2 candidates for subsequent *in vivo* validation.

To further validate LNP mediated antagomir-miR-9-5p delivery in a physiological context, a TBI mouse model was utilized^37, 38^. Mice were subjected to TBI and subsequently treated with LNP41 or LNP12 loaded with antagomir-miR-9-5p *via* intranasal administration. At 24 hours post-injury, RT-qPCR analysis of brain tissue showed that TBI induction sharply upregulated miR-9-5p and inflammatory cytokines. However, intranasal delivery of both LNP41 and LNP12 effectively reversed this pathological alteration. Consistent with the *in vitro* findings, both formulations significantly reduced cerebral miR-9-5p levels and attenuated the expression of TNF-α and IL-1β (Fig. 4D). Based on its consistent superior performance across both transfection efficiency and anti-inflammatory potency, LNP41 was identified as the lead formulation for subsequent long-term therapeutic evaluation.

### Intranasal delivery of miR-9-5p antagomir-loaded LNP-41 promotes functional recovery and modulates chronic neuroinflammation

To evaluate whether the initial suppression of neuroinflammation leads to sustained neurological benefits, the long-term therapeutic efficacy of the lead formulation, LNP41, was assessed in the TBI mouse model over a 28-day period (Fig. 5A). Following TBI induction, mice received three intranasal doses of miR-9-5p antagomir-loaded LNP41 on days 0, 1, and 2. Neurological function was monitored using the modified Neurological Severity Score (mNSS). Untreated TBI mice exhibited severe neurological deficits. High mNSS scores were recorded across motor, balance, and reflex tests. In contrast, mice treated with LNP41 showed a rapid and sustained functional recovery. Significant improvements in neurological scores were observed as early as day 1 post-injury and persisted through day 7 (Fig. 5B).

**Figure 5.**
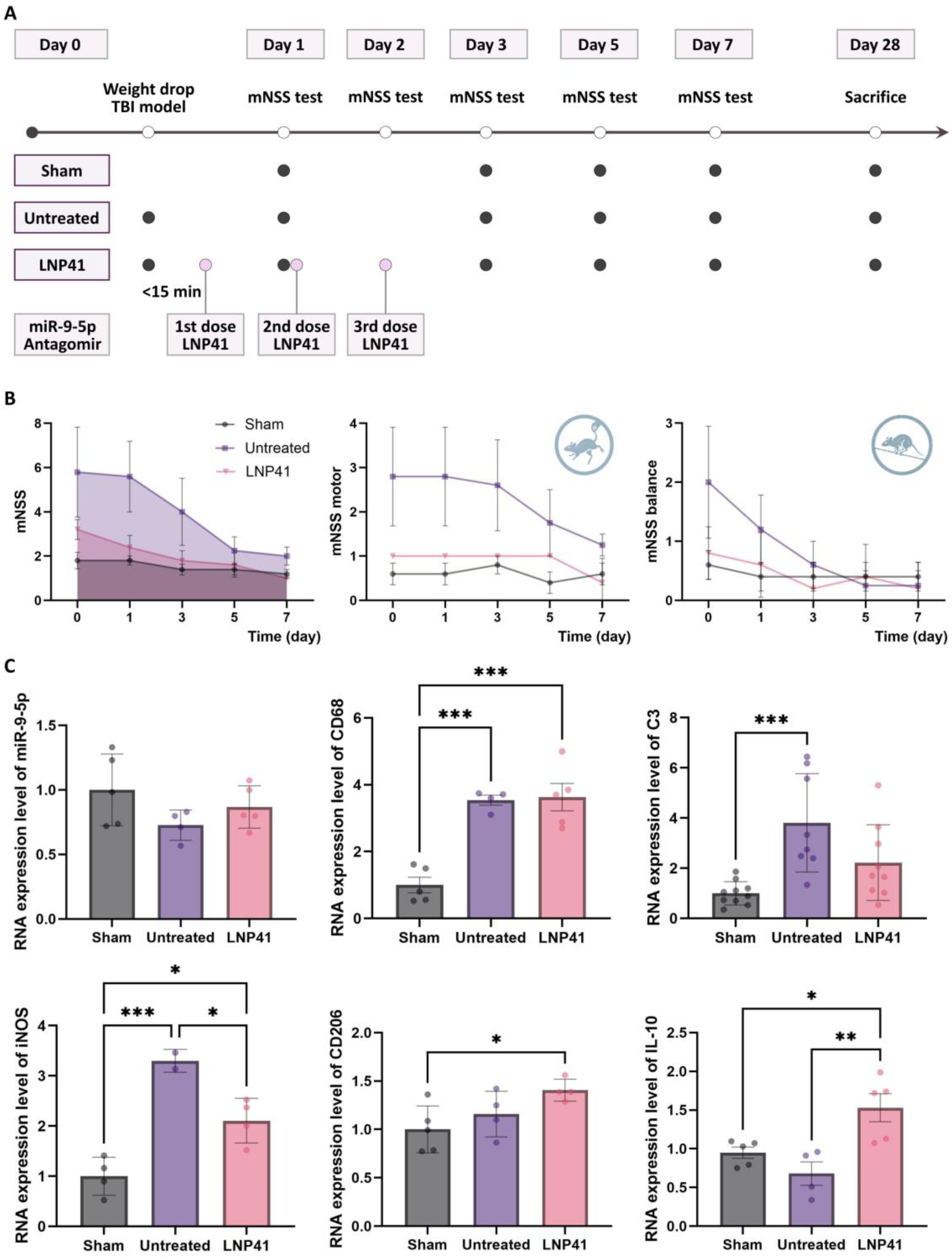
Long-term therapeutic efficacy of LNP41 in a TBI mouse model. **A** Experimental timeline. Mice were assigned to sham, untreated, or treatment groups which received three intranasal doses of LNP41 on days 0, 1, and 2. mNSS was assessed on Days 1, 3, 5, and 7, and tissues were collected on Day 28. **B** Evaluation of neurological function via mNSS (Total, Motor, and Balance scores) from Day 0 to Day 7. LNP41 treatment significantly improved functional outcomes compared to the untreated group. **C** RT-qPCR analysis of gene expression in brain tissue on Day 28. **miR-9-5p:** No significant differences were observed among groups. **CD68:** Significantly upregulated in both untreated and LNP41 groups compared to the sham group. **C3:** Significantly upregulated in the untreated group, while the LNP41 group showed no significant difference compared to the sham group. **iNOS:** Significantly upregulated in the untreated group compared to the sham group. LNP41 treatment significantly reduced iNOS expression compared to the untreated group. **CD206:** Slightly increased in the untreated group compared to the sham group. The LNP41 group showed significantly higher CD206 expression compared to the sham group. **IL-10:** Reduced in the untreated group compared to the sham group. LNP41 treatment significantly increased IL-10 expression compared to the untreated group. Data are presented as mean ± SD. Statistical significance: * *p* < 0.05, ** *p* < 0.01, *** *p* < 0.001, **** *p* < 0.0001.

To investigate the molecular alterations in the chronic phase of TBI, we analyzed gene expression in brain tissues at day 28 post-injury (Fig. 5C). By this late time point, the expression levels of miR-9-5p showed no statistically significant differences among the sham group, untreated and LNP41 treated groups, suggesting that the acute upregulation of miR-9-5p resolves over time. However, markers of chronic neuroinflammation showed distinct patterns. The expression of CD68, a marker of microglial activation and phagocytosis^39, 40^, remained significantly upregulated in both the untreated and LNP41 treated groups compared to sham group, suggesting persistent microglial engagement in the chronic phase. Conversely, complement component C3, a driver of neurotoxicity and synaptic loss^41, 42^, was significantly upregulated in the untreated group but LNP41 treatment effectively mitigated this pathological rise. Collectively, these results suggest that early intranasal intervention with LNP41 confers lasting behavioral protection and selectively modulates specific inflammatory pathways (such as C3-mediated complement activation) in the chronic post-injury brain.

### LNP41 mediated delivery of miRNA-9-5p antagomir attenuates astrocytic C3 expression and restores microglial ramification in the chronic phase of TBI

To further characterize the cellular mechanisms underlying the observed functional recovery, immunofluorescence staining was performed on brain sections collected 28 days post-injury. The analysis specifically focused on two key drivers of chronic neuroinflammation: the upregulation of complement component 3 (C3) in astrocytes and the activation state of microglia.

First, astrocytic responses were assessed by co-staining for GFAP (astrocyte marker) and C3 (Fig. 6A). Untreated TBI mice showed strong C3 expression in GFAP-positive cells. In contrast, intranasal administration of LNP41 markedly reduced astrocytic C3 expression to levels comparable to the sham group (Fig. 6C). This suggests that LNP41 effectively suppresses astrocytic responses associated with C3 expression and A1-like reactivity.

**Figure 6.**
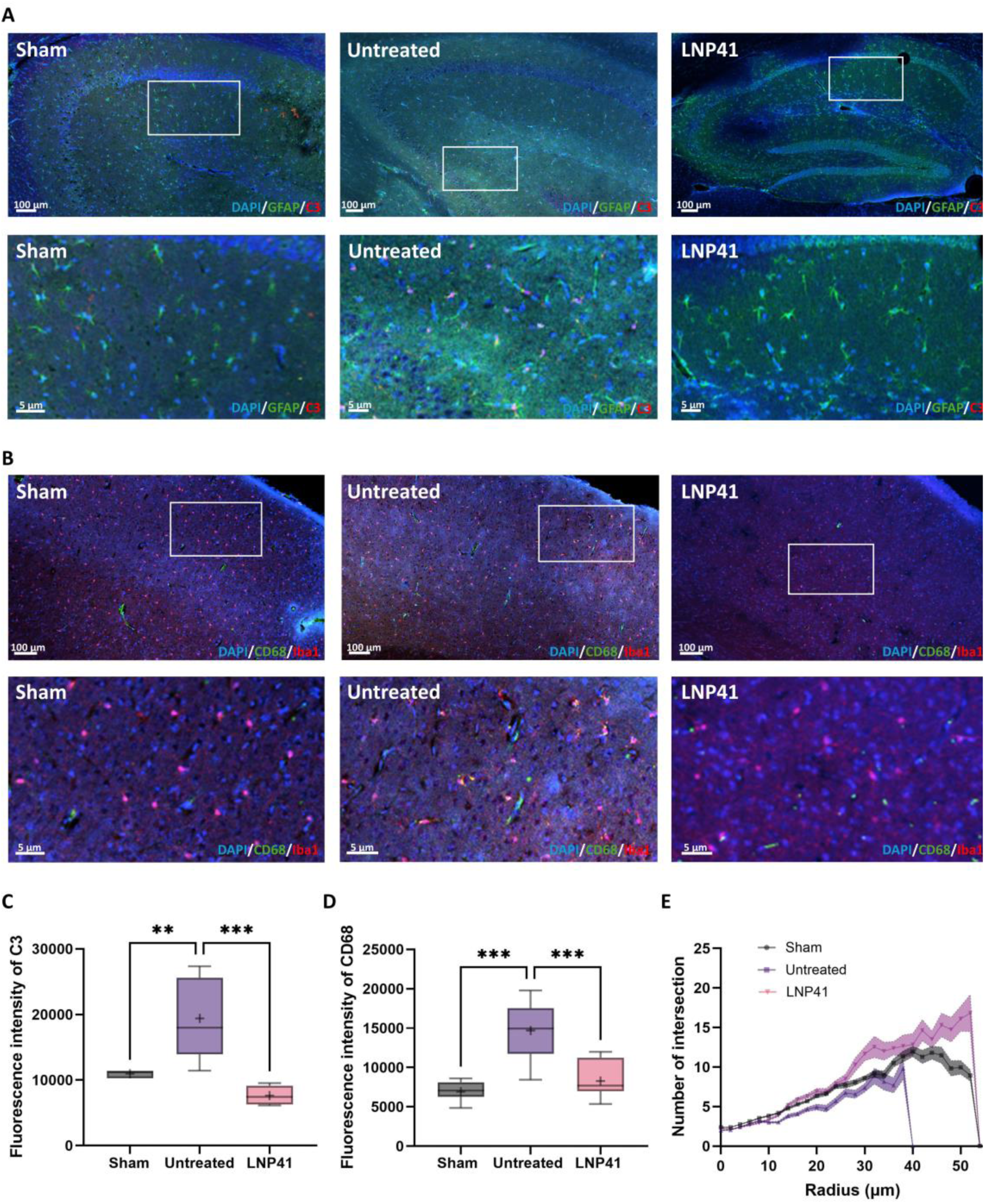
Immunofluorescence analysis of microglia activation and neuroinflammation at 28 days post-TBI. **A** Representative immunofluorescence images of brain sections stained for DAPI (blue), GFAP (green, astrocytes), and C3 (red). Scale bars: 1000 µm (low mag) and 100 µm (high mag). **B** Representative immunofluorescence images stained for DAPI (blue), CD68 (green, activated microglia), and Iba1 (red, microglia). Scale bars: 100 µm (low mag) and 5 µm (high mag). **C** Quantitative analysis of C3 fluorescence intensity. LNP41 treatment significantly reduced astrocytic C3 expression compared to the untreated group. **D** Quantitative analysis of CD68 fluorescence intensity. LNP41 treatment significantly attenuated microglial activation compared to the untreated group. **E** Sholl analysis of microglial morphology showing the number of intersections as a function of radial distance from the soma. LNP41-treated microglia exhibited increased branching complexity compared to the untreated group, with profiles approaching those observed in the sham group. Data are presented as mean ± SD. Statistical significance: * *p* < 0.05, ** *p* < 0.01, *** *p* < 0.001, **** *p* < 0.0001.

Next, microglial activation was examined using Iba1 (microglia marker) and CD68 (lysosomal/phagocytic marker) (Fig. 6B). In the untreated group, microglia exhibited a characteristic amoeboid morphology indicative of activation, co-localizing with high levels of CD68. LNP41 treatment significantly dampened this activation. Quantification revealed a significant reduction in CD68 fluorescence intensity in the LNP41 treated group compared to the untreated group (Fig. 6D).

To provide a more granular assessment of microglial health, Sholl analysis was performed to quantify the morphological complexity of Iba1^+^ cells (Fig. 6E). TBI induction in untreated group led to a drastic reduction in process ramification, resulting in the lowest number of intersections (indicating cell retraction and activation). While LNP41 treatment slightly promoted the formation of highly branched, surveillance-state microglia. The complexity profile showed that microglial complexity was reduced in the untreated group, while LNP41 treatment increased complexity to levels comparable to those observed in the sham group.

Collectively, these histological findings demonstrate that LNP41 treatment profoundly alters the glial microenvironment, suppressing inflammatory markers and restoring healthy cellular morphology.

### Restoration of microglial homeostasis in the olfactory bulb

Given that the intranasal route relies on transport through the olfactory epithelium and trigeminal nerves, we specifically investigated the neuroinflammatory status within the olfactory bulb, the primary entry gateway to the CNS (Fig. 7A).

**Figure 7.**
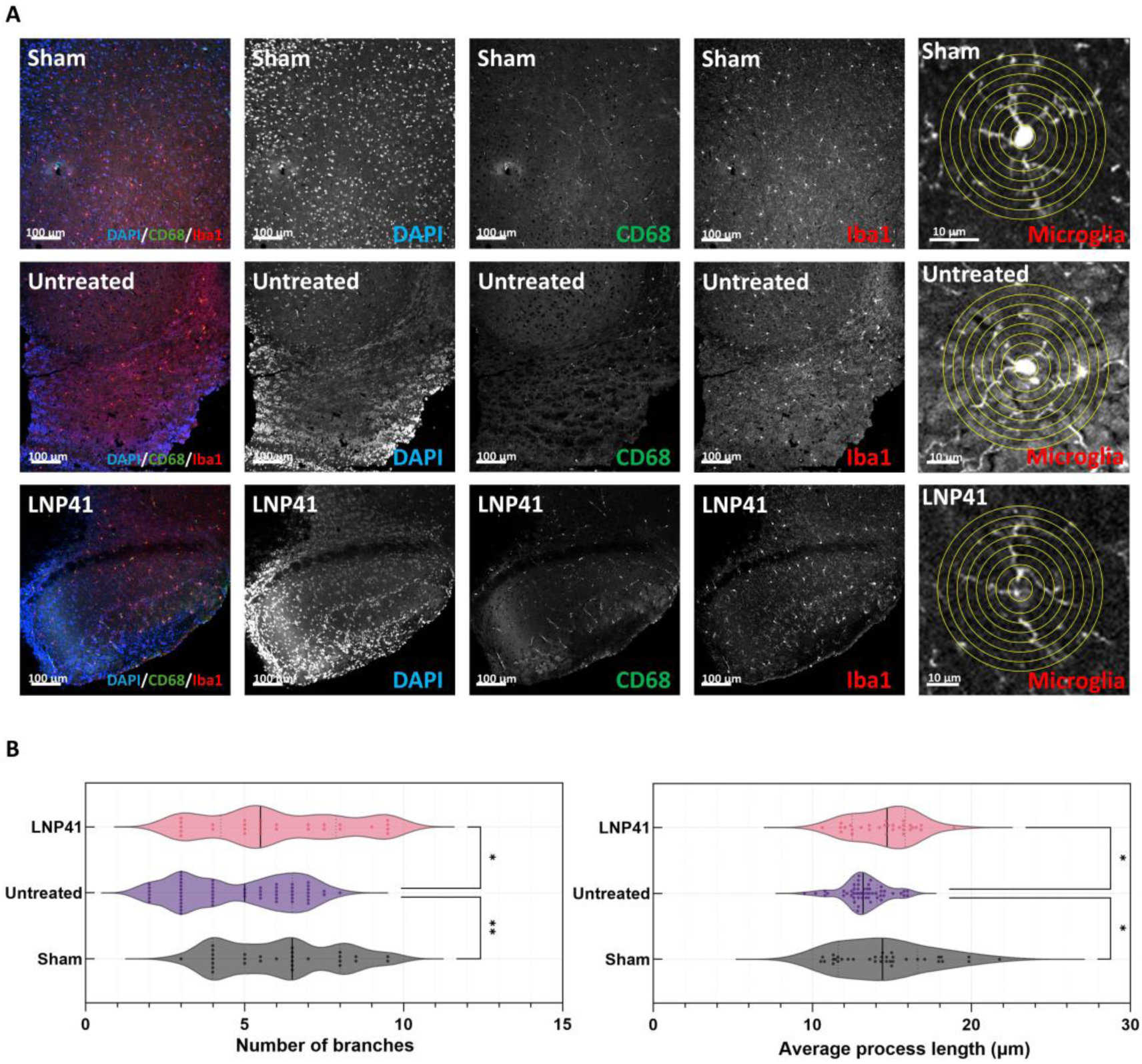
LNP41 treatment restores microglial morphology in the olfactory bulb. **A** Representative immunofluorescence images of the olfactory bulb stained for DAPI (blue), CD68 (green), and Iba1 (red). The rightmost column shows high-magnification images of single microglia with concentric circles illustrating the Sholl analysis strategy. The untreated group induces an amoeboid phenotype, while the LNP41 treated group preserves the ramified state. **B** Quantitative analysis of the number of branches per microglia. The LNP41 treated group shows significantly higher branching complexity compared to the untreated group. **C** Quantitative analysis of the average process length. LNP41 treatment significantly restores process length, preventing the retraction observed in the untreated group. Data are presented as violin plots showing the distribution of individual cells. Statistical significance: * *p* < 0.05, ** *p* < 0.01, *** *p* < 0.001, **** *p* < 0.0001.

Immunofluorescence staining for Iba1 and CD68 revealed distinct morphological alterations in this region. In the untreated group, microglia in the olfactory bulb exhibited a pronounced activated phenotype, characterized by amoeboid cell bodies with retracted, thickened processes and intense CD68 expression. Conversely, treatment with LNP41 effectively preserved the quiescent, ramified morphology of microglia, resembling that observed in the sham group.

To quantitatively assess these morphological changes, a detailed skeletal analysis of individual microglia was performed (Fig. 7A, right panel). Two key metrics of microglial ramification were quantified: the number of branches and the average process length. As illustrated in the violin plots, TBI induction significantly reduced both the complexity and extent of microglial processes compared to the sham group. However, LNP41 treatment effectively reversed this phenotype. The LNP41 treated group exhibited a significantly higher number of branches (Fig. 7B) and increased average process length (Fig. 7C) relative to the untreated group. These findings indicate that intranasal delivery of LNP41 not only mitigates inflammation at the lesion site but also maintains microglial homeostasis along the delivery pathway in the olfactory bulb.

## Discussion

Traumatic brain injury remains a major therapeutic challenge due to limited brain access and persistent neuroinflammation. Here, we developed a DNA barcoded lipid nanoparticle library for intranasal RNA delivery, and identified LNP 41 as a lead candidate enabling efficient brain targeting with minimal peripheral accumulation. Antagomir-miR-9-5p delivery reduced neuroinflammation and improved neurofunctional outcomes, establishing a robust paradigm for non-invasive RNA therapeutics.

The performance of LNP-41 is likely driven by its integrated physiochemical properties rather than any single lipid component. Across formulations, variation in helper lipids led to pronounced differences in normalized brain delivery, with DSPS-containing LNPs showing significantly enhanced delivery (Fig. 4D), indicating a dominant role of helper lipid identity in modulating intranasal transport. This observation aligns with previous studies suggesting that reduced electrostatic mucoadhesion can enhance penetration through the mucus barrier ^19, 43, 44^. In contrast, variation in PEG-lipid terminal functional groups did not produce consistent differences across formulations (Supplementary Fig. E). However, within DSPS-containing LNPs, DSPE-PEG-maleimide consistently ranked among the top-performing combinations, suggesting a context-dependent contribution of PEG functionality (Fig. 4E). Maleimide-modified PEG lipids have been reported to influence protein interactions and cellular association through thiol-reactive chemistry^45^, while phosphatidylserine is known to participate in membrane recognition and uptake processes^46^. Together, these complementary features may contribute to the enhanced normalized brain delivery observed for this formulation, warranting further investigation into the underlying mechanisms.

Building on these findings, intranasal delivery provides a non-invasive route for brain-targeted RNA therapeutics while minimizing peripheral exposure. Systemic LNP administration is often associated with off-target accumulation in peripheral organs, particularly the liver^20, 47^. In contrast, intranasal delivery reduces peripheral exposure by bypassing first-pass hepatic clearance^48, 49^. Our biodistribution data support this distinction, showing minimal accumulation in peripheral organs and strong enrichment in the brain (Fig. 3B). Within the brain, distribution remained spatially heterogeneous. Higher accumulation was observed in regions proximal to the olfactory bulb, with lower levels in deeper parenchymal regions (Supplementary Fig. C), consistent with previous observations^14, 50^. Despite this spatial gradient, robust therapeutic effects were achieved. This suggests that efficacy is not solely determined by bulk nanoparticle accumulation, but also by the biological impact of the delivered cargo. In this context, antagomir-miR-9-5p modulates neuroinflammatory pathways by regulating multiple downstream targets involved in microglial activation^25, 36, 51^, enabling modest delivery to produce amplified biological effects. Consistent with this, *in vitro* inhibition of miR-9-5p suppressed TNF-α and IL-1β genes expression in LPS-challenged microglia (Fig.4C), potentially via attenuation of NF-κB signaling^34^.

*In vivo*, cytokine dynamics showed no significant differences at early time points (Supplementary Fig.F), but diverged at 24 hours with LNP41 treatment attenuating the elevation of pro-inflammatory cytokines (Fig. 5C). Although cytokine levels normalized at later stages, markers of chronic neuroinflammation remained elevated in untreated animals. CD68 protein remained increased (Fig. 6D) despite comparable mRNA levels (Fig. 5C), indicating persistent lysosomal in chronically activated microglia^52, 53^. In parallel, complement component C3 remained elevated in untreated mice, consistent with activation of the astrocyte-microglia axis that drives synaptic pruning and chronic neurodegeneration long after the initial injury^54, 55, 56^. Morphological analysis further supported modulation of microglial activation states. Sholl analysis revealed that microglia in the untreated group retained an amoeboid morphology with reduced process complexity, whereas LNP41 treatment restored a more ramified architecture (Fig.7A). Quantitively, branch complexity increased following treatment and approached levels observed in the sham group (Fig.6E), consistent with a shift toward a less activated microglial phenotype^57, 58^.

Beyond immediate therapeutic efficacy, this study suggests a conceptual shift in nanoparticle design for intranasal delivery. Rather than prioritizing mucoadhesive interactions, which may enhance retention but hinders transport, our findings support lipid compositions that reduce mucosal interactions and facilitate brain access.

Several limitations should be acknowledged. Anatomical differences between rodent and human olfactory systems may influence absolute transport efficiency^59^, warranting validation in higher-order models. In addition, brain distribution showed a spatial gradient, with reduced penetration to deeper brain regions, suggesting that strategies to enhance intracerebral transport may further improve functional outcomes.

In summary, this work establishes an effective strategy framework for intranasal RNA delivery and demonstrates that targeting miR-9-5p can attenuate neuroinflammation following TBI. While the present study focuses on miR-9-5p, the platform is readily adaptable to other RNA cargos and disease contexts. These findings provide a versatile platform that opens new avenues for nose-to-brain delivery across a broad spectrum of central nervous system pathologies.

## Methods

### Materials and animals

#### Materials

The lipid components used for LNP formulation included the ionizable lipid SM102, cholesterol, helper lipids, and PEG-lipids (Fig. 2A, Supplementary Table. 1). The helper lipid library consisted of 7 variants: 3 positively charged species DSTAP (Kyfora Bio, USA), DSPE (BroadPharm, USA), DSPE-Thiol (BroadPharm, USA); 3 negatively charged lipids DSPS (BroadPharm, USA), DSPE-succinic acid (BroadPharm, USA), DSPS-PEG-NHS 600 (BroadPharm, USA); and 1 neutral control DSPC (BroadPharm, USA). The PEG-lipid library included 4 functionalized variants with reactive headgroups DSPE-PEG-Ald 2000 (Nanosoft Polymers, USA), DSPE-PEG-Maleimide 2000 (BroadPharm, USA), DSPE-PEG-COOH 2000 (BroadPharm, USA), DSPE-PEG-NH2 2000 (Avanti Polar Lipids, USA) and DMG-PEG 2000 (Avanti Polar Lipids, USA) as a standard control. DNA barcodes (61 nucleotides with unique 8-nt sequences) were synthesized by Integrated DNA Technologies (IDT, USA). MiR-9-5p antagomir was using miRIDIAN microRNA Mouse mmu-miR-9-5p inhibitor (Horizon Discovery, USA)

#### Animals

Female C57BL/6J mice (4–6 weeks, 18–25 g) were housed under standard conditions with free access to food and water. All animal procedures were approved by the Institutional Animal Care and Use Committee (IACUC) of the University of Massachusetts Amherst (Protocol No.5234).

### Construction and characterization of DNA-barcoded LNP library

To optimize intranasal delivery, a library of 103 mucoadhesive LNP formulations was generated by systematically varying the ratios of helper lipids and PEG-lipids. Each LNP formulation was encapsulated with a 61 nucleotides DNA barcode containing 8-nt unique sequence to enable specific identification (Supplementary Table. 2). LNPs were prepared using a Sunshine Prime Microfluidic (Unchained Labs, USA) by mixing the lipid phase with the aqueous phase containing the DNA barcodes. Following formulation, particles were dialyzed against PBS (pH 7.4, 6 kDa cutoff) for 4 hours at 4°C to remove ethanol and unencapsulated cargo. Physicochemical properties, including hydrodynamic size and polydispersity index (PDI), were measured by dynamic light scattering using a Zetasizer ZSP (Malvern Panalytical, UK). Encapsulation efficiency (EE) was assessed using the Quant-iT RiboGreen RNA assay (Thermo Fisher). Formulations meeting the quality control criteria of size <300 nm, PDI <0.4, and EE >50% were selected for subsequent *in vivo* screening.

### High-throughput *in vivo* screening for brain distribution

#### Preparation for pooled library

To identify formulations with optimal brain-targeting capability, a pooled library containing equimolar amounts of all qualified barcoded LNPs was prepared. The pooled library was intranasally administered to C57BL/6J mice (5-6 weeks) via a refined catheter-directed olfactory delivery method ^12, 35^.

#### Olfactory region-specific intranasal administration

Briefly, mice were anesthetized with isoflurane and placed in a supine position with the head elevated at approximately 45° relative to the body to facilitate access to the nostril. The head was gently stabilized using one hand, while the catheter was held in the other hand. Nutriline Twinflo catheters (outer diameter ∼0.6 mm, Vygon, France) were trimmed to a length of approximately 3 cm. The lateral surface of the distal tip was coated with a thin layer of petroleum jelly (Vaseline) to reduce mucosal friction and prevent spreading of the formulation to the lower respiratory region. The catheter was introduced into one nostril at a minimal insertion angle of approximately 20-25° relative to the nasal line to ensure targeting of the dorsal meatus. During advancement, the catheter was gently rotated to navigate the narrow nasal passage. The catheter was advanced until slight resistance was encountered at the level of the ethmoid turbinate, corresponding to the olfactory region. In C57BL/6 mice, the mean insertion depth for olfactory targeting is approximately 10-11 mm, depending on body weight. The catheter was not advanced beyond this point to avoid mucosal injury or misplacement into the nasopharynx.

After correct positioning, the formulation was infused using a 10 µL Microliter Syringe (Model 701 RN, 26s gauge, 2 in, Hamilton Company, USA). The catheter was then carefully withdrawn while maintaining the animal in a supine position for approximately one minute to minimize drainage. To achieve a total dosage of 20 µL (0.02 mg/kg) while minimizing fluid runoff, the administration was split into two 10 µL doses separated by an interval of at least 15 minutes.

#### Tissue collection and analysis

One hour post-administration, mice were sacrificed, and tissues were collected, including brain subregions (e.g., olfactory bulb, cortex), liver, spleen, lung, heart, and kidney. For cell-type specific analysis, neurons, microglia, astrocytes, and oligodendrocytes were isolated from brain tissue using a neuron isolation kit and specific microbeads (Miltenyi Biotec, Germany). Tissue and cell samples were homogenized, and total DNA was extracted using the QuickExtract DNA Extraction Solution (Biosearch Technologies, USA). Barcode sequences were amplified using Illumina-compatible primers and sequenced via the MiSeq platform at the Genomics Core Facility, Harvard Medical School. The relative abundance of each barcode in the collected tissues was quantified and processed using a two-step normalization workflow. First, normalized delivery was calculated by normalizing barcode counts within each sample to total counts, internal controls, and the input LNP library, followed by averaging across biological replicates. Second, accumulation was derived by scaling normalized delivery values to qPCR-measured DNA levels and further normalizing to the lowest signal within each tissue or cell-type group. These metrics were used to generate biodistribution heatmaps and identify top-performing LNPs for further validation.

### Stepwise i*n vitro* validation

#### Cell culture

BV2 murine microglial cells (ATCC, USA) were cultured in high-glucose Dulbecco’s Modified Eagle Medium (DMEM, Fisher Scientific, USA) supplemented with 10% heat-inactivated fetal bovine serum (FBS, Sigma-Aldrich, USA) and 1% penicillin-streptomycin (Cytiva, USA). Cells were maintained at 37 °C in a humidified incubator containing 5% CO₂. Passaging was performed at 70-85% confluence using 0.05% trypsin-EDTA (Thermo Fisher Scientific, USA). For functional assays, BV2 cells were seeded into 6-well plates at a density of 5.0×10⁵ cells/well for RT-qPCR analysis, or into 96-well plates at 2.0×10⁴ cells/well for luciferase readouts. All experiments were conducted using cells within 20 passages and confirmed to be in the logarithmic growth phase.

#### Luciferase reporter assay

To evaluate transfection efficiency, CleanCap® M6 FLuc mRNA (Maravai LifeSciences, USA) was encapsulated into the top 10 LNP formulations. BV2 microglial cells were treated with FLuc-LNPs at a dose of 20 ng mRNA/well and incubated for 6 hours. Luminescence intensity was subsequently quantified using the Pierce Firefly Luciferase Glow Assay Kit (Thermo Fisher Scientific, USA) on a BioTek Synergy 2 plate reader (BioTek, Agilent Technologies, USA).

#### miR-9-5p knockdown and cytokine analysis

For gene silencing validation, BV2 cells were transfected with LNPs loaded with antagomir-miR-9-5p (50 nM final concentration) for 24 hours. To induce an inflammatory response, cells were subsequently stimulated with lipopolysaccharides (LPS, Sigma-Aldrich, USA) at 1 µg/mL for 2 hours. Total RNA was extracted using the RNAzol reagent (Sigma-Aldrich, USA) according to the manufacturer’s protocol.

For microRNA quantification, a two-step poly(A)-tailing RT-qPCR method was employed. First, polyadenylation was performed using the Poly(A) Polymerase Tailing Kit (Epicentre, Illumina, USA). cDNA synthesis was then conducted via oligo(dT)-adapter reverse transcription using the GoScript Reverse Transcription System (Promega, USA). qPCR was performed using the Luna Universal qPCR Master Mix (New England Biolabs, USA), with U6 snRNA serving as the internal reference. Primers are listed in Supplementary Table 3.

For pro-inflammatory cytokine analysis, the expression levels of IL-1β and TNF-α were measured using the Luna Universal One-Step RT-qPCR Kit (New England Biolabs, USA). GAPDH was used as internal control for normalization. Relative gene expression was calculated using the 2^-ΔΔCt^ method, and data are presented as mean ± SD.

### *In vivo* validation in TBI mouse model

#### TBI model induction and treatment regimen

Mice were randomly assigned to three experimental groups: Negative control (NC), TBI model (PC), and TBI treated with LNPs loaded with antagomir-miR-9-5p. The TBI model was induced using a weight-drop method^37, 60^. Briefly, a 54 g weight was dropped from a height of 71 cm (28 inches) on the cortical region of the skull to induce traumatic brain injury. Treatments were administered intranasally (20 µL/dose) via catheter-directed delivery. The initial dose was administered 15–30 minutes post-injury.

*For short-term analysis:* Mice were perfused and sacrificed at 6 hours and 24 hours post-injury.

*For long-term analysis:* Mice received a total of three intranasal doses on days 0, 1, and 2 post-injury, then perfused and sacrificed on day 28.

#### Gene expression analysis

For both short-term and long-term studies, brain tissues were collected from the lesion site at 6 hours, 24 hours, and 28 days post-injury. Tissue homogenization was performed, and total RNA was extracted using the RNAzol reagent (Sigma-Aldrich, USA) following the manufacturer’s instructions. The expression levels of miR-9-5p and inflammatory markers (IL-1β, TNF-α, CD68, C3) were quantified by RT-qPCR as described in Section 2.4.3.

#### Neurological function assessment (mNSS)

For long-term studies, neurological function was evaluated on days 0, 1, 3, 5, and 7 post-injury using the modified Neurological Severity Score (mNSS). This composite scoring system assesses motor, sensory, balance, and reflex functions, with scores ranging from 0 (normal function) to 18 (maximal neurological deficit). Detailed scoring criteria are provided in Supplementary Table 4.

#### Immunofluorescence histology

On day 28 post-injury, mice were perfused with PBS followed by 4% paraformaldehyde (PFA). Brains were harvested, post-fixed, and embedded in OCT compound (Sakura Finetek, USA). Frozen sections were cut to a thickness of 30 µm using a cryostat. For immunofluorescence staining, free-floating sections were incubated for 48 hours at 4 °C with primary antibodies against Iba1 (1:500, Abcam, UK), CD68 (1:250, Abcam, UK), GFAP (1:500, Abcam, UK), and C3 (1:250, Abcam, UK). Sections were subsequently incubated with fluorophore-conjugated secondary antibodies and mounted using ProLong Diamond Antifade Mountant with DAPI (Thermo Scientific, USA).

Images were acquired using an AXR-NSPARC confocal microscope (Nikon, Japan). Quantitative analysis of fluorescence intensity and cell morphology was performed using Fiji/ImageJ software (Version 2.14, National Institutes of Health, USA).

### Statistical analysis

All quantitative data are expressed as mean ± standard deviation (SD). Differences among multiple groups were assessed using one-way analysis of variance (ANOVA) followed by Tukey’s post-hoc test. Statistical analyses were performed using GraphPad Prism 11 (GraphPad Software, USA). A *p*-value < 0.05 was considered statistically significant. Significance levels are denoted as follows: * *p* < 0.05, ** *p* < 0.01, *** *p* < 0.001, and **** *p* < 0.0001.

## Data availability statement

All data that support the findings of this study are included within the article (and any supplementary files).

## Acknowledgments

This study was supported by Alzheimer’s Association Research Grants (AARG-25-1487957)

## Author information

These authors contributed equally: Yihan Xing, Vy Do.

## Contributions

J.G. conceived and designed the entire project. Y.X and V.D. performed all of the experiments. V.D., Z.X., and C.D. generated sequencing data. Z.X., C.Y., Z.Z provided methodology. Y.X. wrote the manuscript. All authors reviewed and approved the final version of the manuscript.

## Ethics declarations

All animal experiments were approved by the Institutional Animal Care and Use Committee, University of Massachusetts Amherst, and performed following the animal use protocol (No. 5234).

