## Supplemental Figure for "Combinatorial screening of nanoparticles for nose-to-brain RNA delivery to modulate neuroinflammation after traumatic brain injury"

**Supplementary Information**

**Supplementary Figure: Physicochemical characterization biodistribution and gene expression of LNP formulations.**

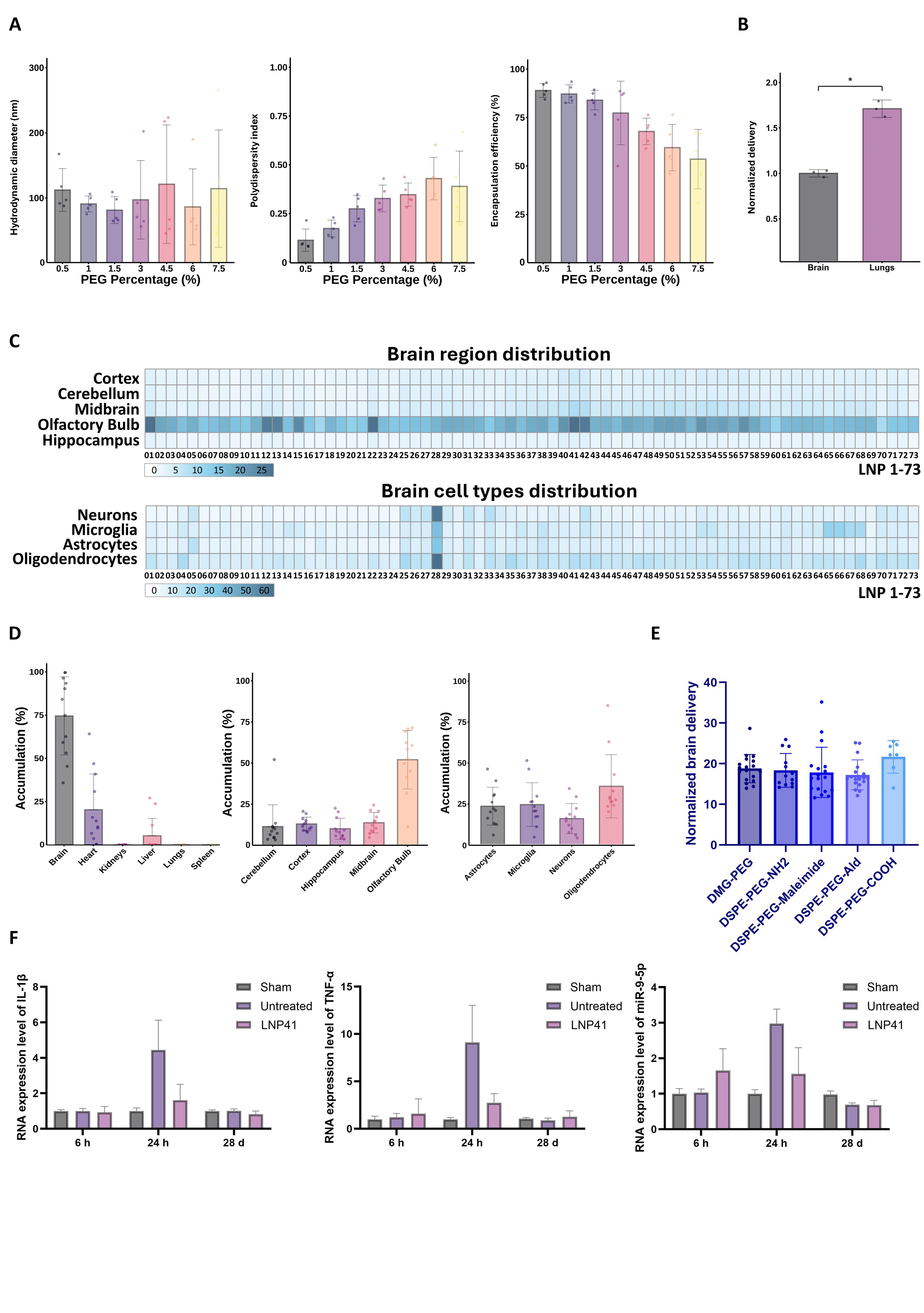

**A** Effects of PEG lipid content on LNP physicochemical properties. Hydrodynamic diameter, polydispersity index, and encapsulation efficiency were measured for LNPs formulated with increasing PEG percentages. Individual data points represent independent formulations, with bars indicating mean values. **B** Normalized delivery of the selected LNP formulation in the brain and lungs following intranasal administration. Data are shown as mean values with statistical comparison as indicated. **C** Heatmaps showing regional brain distribution (top) and brain cell type distribution (bottom) of DNA-barcoded LNPs after intranasal administration. Color intensity represents normalized delivery across brain regions and neural cell populations. **D** Quantification of LNP accumulation across major brain regions (including cortex, cerebellum, hippocampus and olfactory bulb), as well as major brain cell types (including astrocytes, neurons, microglia and oligodendrocytes). **E** Normalized brain delivery of LNPs stratified by PEG-lipid terminal chemistry. No consistent differences were observed across PEG end groups, indicating a limited contribution of PEG functionality as an isolated variable. **F** Temporal expression of inflammatory markers (IL-1β, TNF-α) and miR-9-5p in the TBI model following intranasal administration of the selected antagomir-loaded LNP. RNA levels were measured at 6 h, 24 h, and 28 days post-treatment.
