## Supplemental Table for "Combinatorial screening of nanoparticles for nose-to-brain RNA delivery to modulate neuroinflammation after traumatic brain injury"

**Supplementary Table 1. LNP library formulations**

| **ID** | **Helper lipid** | **Cholesterol %** | **PEG Type** | **PEG %** |
| --- | --- | --- | --- | --- |
| 1 | DSPC | 39.5 | DMG-PEG 2000 | 0.5 |
| 2 | DSPC | 39 | DMG-PEG 2000 | 1 |
| 3 | DSPC | 38.5 | DMG-PEG 2000 | 1.5 |
| 4 | DSPC | 35.5 | DMG-PEG 2000 | 4.5 |
| 5 | DSTAP | 38.5 | DMG-PEG 2000 | 1.5 |
| 6 | DSTAP | 35.5 | DMG-PEG 2000 | 4.5 |
| 7 | DSPE | 38.5 | DMG-PEG 2000 | 1.5 |
| 8 | DSPE | 35.5 | DMG-PEG 2000 | 4.5 |
| 9 | DSPE-Thiol | 38.5 | DMG-PEG 2000 | 1.5 |
| 10 | DSPE-Thiol | 35.5 | DMG-PEG 2000 | 4.5 |
| 11 | DSPE-Thiol | 32.5 | DMG-PEG 2000 | 7.5 |
| 12 | DSPC | 39.5 | DSPE-PEG-NH2 2000 | 0.5 |
| 13 | DSPC | 39 | DSPE-PEG-NH2 2000 | 1 |
| 14 | DSPC | 38.5 | DSPE-PEG-NH2 2000 | 1.5 |
| 15 | DSPC | 37 | DSPE-PEG-NH2 2000 | 3 |
| 16 | DSPC | 35.5 | DSPE-PEG-NH2 2000 | 4.5 |
| 17 | DSPC | 34 | DSPE-PEG-NH2 2000 | 6 |
| 18 | DSTAP | 38.5 | DSPE-PEG-NH2 2000 | 1.5 |
| 19 | DSTAP | 35.5 | DSPE-PEG-NH2 2000 | 4.5 |
| 20 | DSPE | 38.5 | DSPE-PEG-NH2 2000 | 1.5 |
| 21 | DSPE | 35.5 | DSPE-PEG-NH2 2000 | 4.5 |
| 22 | DSPE-Thiol | 38.5 | DSPE-PEG-NH2 2000 | 1.5 |
| 23 | DSPE-Thiol | 35.5 | DSPE-PEG-NH2 2000 | 4.5 |
| 24 | DSPE-Thiol | 32.5 | DSPE-PEG-NH2 2000 | 7.5 |
| 25 | DSPC | 39.5 | DSPE-PEG-Maleimide 2000 | 0.5 |
| 26 | DSPC | 39 | DSPE-PEG-Maleimide 2000 | 1 |
| 27 | DSPC | 38.5 | DSPE-PEG-Maleimide 2000 | 1.5 |
| 28 | DSPC | 37 | DSPE-PEG-Maleimide 2000 | 3 |
| 29 | DSPC | 35.5 | DSPE-PEG-Maleimide 2000 | 4.5 |
| 30 | DSPC | 34 | DSPE-PEG-Maleimide 2000 | 6 |
| 31 | DSPC | 32.5 | DSPE-PEG-Maleimide 2000 | 7.5 |
| 32 | DSTAP | 38.5 | DSPE-PEG-Maleimide 2000 | 1.5 |
| 33 | DSTAP | 35.5 | DSPE-PEG-Maleimide 2000 | 4.5 |
| 34 | DSTAP | 32.5 | DSPE-PEG-Maleimide 2000 | 7.5 |
| 35 | DSPE | 38.5 | DSPE-PEG-Maleimide 2000 | 1.5 |
| 36 | DSPS | 35.5 | DMG-PEG 2000 | 4.5 |
| 37 | DSPE-PEG-NHS 600 | 38.5 | DMG-PEG 2000 | 1.5 |
| 38 | DSPE-PEG-NHS 600 | 35.5 | DMG-PEG 2000 | 4.5 |
| 39 | DSPE-Succinic Acid | 38.5 | DMG-PEG 2000 | 1.5 |
| 40 | DSPE-Succinic Acid | 35.5 | DMG-PEG 2000 | 4.5 |
| 41 | DSPS | 38.5 | DSPE-PEG-Maleimide 2000 | 1.5 |
| 42 | DSPS | 35.5 | DSPE-PEG-Maleimide 2000 | 4.5 |
| 43 | DSPS | 32.5 | DSPE-PEG-Maleimide 2000 | 7.5 |
| 44 | DSPE-PEG-NHS 600 | 35.5 | DSPE-PEG-Maleimide 2000 | 4.5 |
| 45 | DSPE-PEG-NHS 600 | 32.5 | DSPE-PEG-Maleimide 2000 | 7.5 |
| 46 | DSPE-Succinic Acid | 38.5 | DSPE-PEG-Maleimide 2000 | 1.5 |
| 47 | DSPE-Succinic Acid | 35.5 | DSPE-PEG-Maleimide 2000 | 4.5 |
| 48 | DSPS | 35.5 | DSPE-PEG-Ald 2000 | 4.5 |
| 49 | DSPE-PEG-NHS 600 | 38.5 | DSPE-PEG-Ald 2000 | 1.5 |
| 50 | DSPE-PEG-NHS 600 | 35.5 | DSPE-PEG-Ald 2000 | 4.5 |
| 51 | DSPE-Succinic Acid | 38.5 | DSPE-PEG-Ald 2000 | 1.5 |
| 52 | DSPE-Succinic Acid | 35.5 | DSPE-PEG-Ald 2000 | 4.5 |
| 53 | DSPE-Succinic Acid | 32.5 | DSPE-PEG-Ald 2000 | 7.5 |
| 54 | DSPE-PEG-NHS 600 | 38.5 | DSPE-PEG-COOH 2000 | 1.5 |
| 55 | DSPE-Succinic Acid | 35.5 | DSPE-PEG-COOH 2000 | 4.5 |
| 56 | DSPE-Succinic Acid | 32.5 | DSPE-PEG-COOH 2000 | 7.5 |
| 57 | DSPC | 39.5 | DSPE-PEG-COOH 2000 | 0.5 |
| 58 | DSPC | 39 | DSPE-PEG-COOH 2000 | 1 |
| 59 | DSPC | 38.5 | DSPE-PEG-COOH 2000 | 1.5 |
| 60 | DSPC | 37 | DSPE-PEG-COOH 2000 | 3 |
| 61 | DSPC | 39.5 | DSPE-PEG-Ald 2000 | 0.5 |
| 62 | DSPC | 39 | DSPE-PEG-Ald 2000 | 1 |
| 63 | DSPC | 38.5 | DSPE-PEG-Ald 2000 | 1.5 |
| 64 | DSPC | 37 | DSPE-PEG-Ald 2000 | 3 |
| 65 | DSPC | 35.5 | DSPE-PEG-Ald 2000 | 4.5 |
| 66 | DSPC | 34 | DSPE-PEG-Ald 2000 | 6 |
| 67 | DSPC | 32.5 | DSPE-PEG-Ald 2000 | 7.5 |
| 68 | DSPE-Thiol | 38.5 | DSPE-PEG-Ald 2000 | 1.5 |
| 69 | DSPE-Thiol | 35.5 | DSPE-PEG-Ald 2000 | 4.5 |
| 70 | DSTAP | 38.5 | DSPE-PEG-Ald 2000 | 1.5 |
| 71 | DSTAP | 35.5 | DSPE-PEG-Ald 2000 | 4.5 |
| 72 | DSTAP | 32.5 | DSPE-PEG-Ald 2000 | 7.5 |
| 73 | DSPE | 32.5 | DMG-PEG 2000 | 6 |
| Percentages shown in this table represent molar percentages (mol%) of the total lipid composition. The ionizable lipid component was fixed at 50% SM-102. Helper lipids were maintained at 10 mol% of the total lipid content. Cholesterol was used in its unmodified form and kept constant across all formulations. | | | | |

**Supplementary Table 2. DNA barcodes list**

| **ID** | **Unique 8-nt Sequence (5’-3’)** | **ID** | **Unique 8-nt Sequence (5’-3’)** |
| --- | --- | --- | --- |
| 1 | TGA TAT TG | 26 | TCC TAA GA |
| 2 | GAC GCA AT | 27 | CAA GAA GG |
| 3 | GCG AGT AT | 28 | TAG AAT TA |
| 4 | ACC TAA TC | 29 | GGC GCC AA |
| 5 | AGG CGC TA | 30 | TAG ATC CG |
| 6 | GAT CTA CC | 31 | CGA GCA GC |
| 7 | CTA CTG AT | 32 | TAA GAT GA |
| 8 | TGA TCT AT | 33 | AGC TCG GA |
| 9 | ATG AGA TG | 34 | TAA CCG AA |
| 10 | GCG AAT TC | 35 | TAT ATC TA |
| 11 | GAT TCC GG | 36 | AAG AGG AT |
| 12 | ATA ATA TA | 37 | ACG TCG AA |
| 13 | AGC ATG CG | 38 | CAT CAT TA |
| 14 | GAT TCA AC | 39 | TTG CAA CT |
| 15 | TAC CTG CT | 40 | TCT AAC TG |
| 16 | GCT AAT CG | 41 | TAT GCC TT |
| 17 | CTC CTT CG | 42 | GTA ATT GC |
| 18 | ACG CTA GC | 43 | GTC TCC GT |
| 19 | GCA GGA CT | 44 | TGC ATG GT |
| 20 | ATT GCT CT | 45 | AGT CCG GT |
| 21 | TAC GCT CG | 46 | TCC TGA TG |
| 22 | ACG CTC CA | 47 | ATC GTC TA |
| 23 | CGG TCA AT | 48 | GGA CGT CC |
| 24 | CGC CTA TT | 49 | CTA CGA GG |
| 25 | TTG CGT TG | 50 | CAA TCC GT |
| Only the unique 8-nt barcode regions are listed. All barcodes were inserted into the following oligonucleotide scaffold: 5’ GAT* GCA CGC CTT ACG ACT CAT CTN WNH [8-nt barcode] N WHG TGG TTA GTC GAG CAG AGA CTA* G 3’ Degenerate bases follow IUPAC nomenclature. Asterisks (*) indicate phosphorothioate linkages. | | | |

**Supplementary Table 3. Primer list**

| **Gene** | **Primer （5’-3’）** |
| --- | --- |
| *Gadph* | Forward: TCAGCAATGCCTCCTGCAC  Reverse: TCTGGGTGGCAGTGATGGC |
| *Tnfα* | Forward: CTGAACTTCGGGGTGATCGG  Reverse: GGCTTGTCACTCGAATTTTGAGA |
| *Il1β* | Forward: TGGAGAGTGTGGATCCCAAG  Reverse: GGTGCTGATGTACCAGTTGG |
| *Cd68* | Forward: TGTCTGATCTTGCTAGGACCG  Reverse: GAGAGTAACGGCCTTTTTGTGA |
| *C3* | Forward: CCAGCTCCCCATTAGCTCTG  Reverse: GCACTTGCCTCTTTAGGAAGTC |
| *Inos* | Forward: CAGGGAGAACAGTACATGAACAC  Reverse: TTGGATACACTGCTACAGGGA |
| *CD206* | Forward: AGACGAAATCCCTGCTACTG  Reverse: CACCCATTCGAAGGCATTC |
| *IL-10* | Forward: GCCAGAGCCACATGCTCCTA  Reverse: GATAAGGCTTGGCAACCCAAGTAA |
| *U6* | Forward: CTCGCTTCGGCAGCACA  Reverse: TCCGATCGTGAAGCGTTC |
| *Mir-9-5p* | Forward: TCATACAGCTAGATAACCAAAGA  Reverse: TCTTTGGTTATCTAGCTGTATGA |

**Supplementary Table 4. mNSS tests and scoring values**

| **Category** | **Test item** | **Criteria** | **Score** |
| --- | --- | --- | --- |
| **Motor test**  Max score 6 | Forelimb flexion | Flexion of forelimb upon lifting | 1 |
|  | Hindlimb flexion | Flexion of hindlimb upon lifting | 1 |
|  | Head deviation | Head deviates >10° to vertical axis within 30s | 1 |
|  | Walking test | Normal walk | 0 |
|  |  | Inability to walk straight | 1 |
|  |  | Circling toward paretic side | 2 |
|  |  | Falls down to paretic side | 3 |
| **Sensory test**  Max score 2 | Placing test | Visual and tactile test (observe reaching response) | 0 |
|  |  | Absence of response | 1 |
|  | Proprioceptive test | Deep sensation (push paw against table to stimulate muscles) | 0 |
|  |  | Absence of response (no resistance/adjustment) | 1 |
| **Beam balance**  Max score 6 | Beam test | Balances with steady posture | 0 |
|  |  | Grasps side of beam | 1 |
|  |  | Hugs beam and 1 limb falls down | 2 |
|  |  | Hugs beam and 2 limbs fall down, or spins (>60s) | 3 |
|  |  | Attempts to balance, but falls off (>40 s) | 4 |
|  |  | Attempts to balance, but falls off (>20 s) | 5 |
|  |  | Falls off; no attempt to balance or hang on (<20 s) | 6 |
| **Reflexes**  Max score 4 | Pinna reflex | Head shakes when auditory meatus is touched | 0 |
|  |  | Absence of reflex | 1 |
|  | Corneal reflex | Eye blinks when cornea is lightly touched | 0 |
|  |  | Absence of reflex | 1 |
|  | Startle reflex | Motor response to a brief noise (e.g., snapping paper) | 0 |
|  |  | Absence of reflex | 1 |
|  | Abnormal movements | Seizures, myoclonus, dystonia, etc. | 0 |
|  |  | Presence of abnormal movements | 1 |
| Score value of 1 was given for the inability to perform a test, or for the lack of a tested reflex,  or for abnormal movement^61, 62^. | | | |
